# Stable long-term individual variation in chimpanzee technological efficiency

**DOI:** 10.1101/2023.11.21.568000

**Authors:** S. Berdugo, E. Cohen, A. J. Davis, T. Matsuzawa, S. Carvalho

**Affiliations:** Social Body Lab, Institute of Human Sciences, University of Oxford, Oxford; Primate Models for Behavioural Evolution Lab, Institute of Human Sciences, University of Oxford, Oxford; Wadham College, Parks Road, Oxford, OX1 3PN, United Kingdom; Chubu Gakuin University, Gifu, Japan; Division of the Humanities and Social Sciences, California Institute of Technology, Pasadena, CA, USA; College of Life Science, Northwest University, Xi’an 710069, China; ICArEHB, Interdisciplinary Center for Archaeology and Evolution of Human Behaviour FCHS, Universidade do Algarve, Faro 8005, Portugal; Gorongosa National Park, Sofala, Mozambique

## Abstract

Using tools to access hard-to-reach and high-quality resources, such as termites, honey, and nuts, initiated a fundamental adaptive shift in human and nonhuman primate cognitive and behavioural evolution. Variation in the efficiency of extracting calorie-rich and nutrient-dense resources directly impacts energy expenditure, and potentially has significant repercussions for cultural transmission where model selection biases are employed during skill acquisition. Assessing variation in efficiency is key to understanding the evolution of complex behavioural traits in primates. Yet, individual-level differences beyond age- and sex-class in primate extractive foraging efficiency have never been investigated. Here, we used 25 years (1992– 2017) of video data of the Bossou chimpanzee community (Guinea), to investigate whether individual differences in nut-cracking efficiency exist across the life span of chimpanzees aged ≥ 6 years. Data from 3,882 oil-palm nut-cracking bouts from over 800 hours of observation were collected. We found long-term stable and reliable individual differences in four (out of five) measures of efficiency. We found no sex effect, challenging previous research on a female bias in chimpanzee tool use. These life-long differences in extractive foraging impacts daily energy budgets, which potentially have significant individual fitness and life history consequences. Additionally, the establishment of long-term individual variation in chimpanzee stone tool use has implications for interpreting archaeological records of hominins. Our findings highlight the importance of longitudinal data from long-term field sites when investigating underlying cognitive and behavioural diversity across individual lifespans and between populations.

## Main

The importance of individual variation in cognition and behaviour is increasingly appreciated in research on nonhuman animals^1^. Such variation has ramifications at both the individual and population levels, with broader implications for life history^2–4^, cultural evolution, and interpretations of the archaeological record^5^ (see Figure 1). For example, a recent large-scale meta-analysis found individual variation in the migration timing of land-, water-, and seabirds, which impacts the breeding success and survival of a given migration^6^.

**Figure 1.**
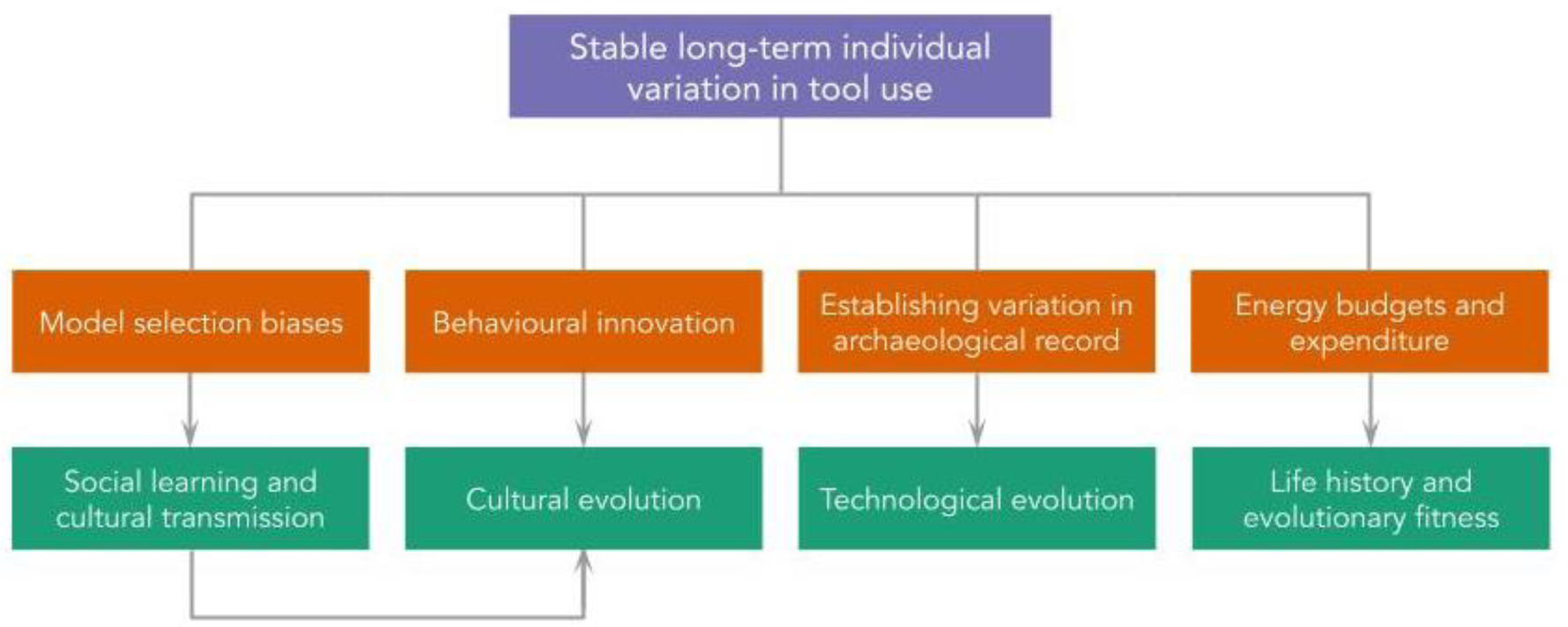
Implications of long term stable individual variation in lithic technological behaviour.

Yet, although cognitive and behavioural variation has been reported, the research has mostly focused on one time-point for each individual or is limited to short time spans^1,7,8^. We argue that this approach does not reflect the true extent and patterns of variation within wild populations. Here, we aim to fill this knowledge gap and establish for the first time long-term stable and reliable individual differences in chimpanzee technological efficiency.

Firstly, persistent variation in the proficiency and efficiency with which individuals extract high quality resources from the environment directly impacts energy expenditure and individual fitness. An individual who is slow or inefficient at a given extractive-foraging technique relative to others in the population will have less time and energy for other fitness-enhancing tasks, thus incurring relative fitness costs^9^. This is particularly true for complex tool-assisted foraging tasks aimed at extracting high calorie resources, such as wood-boring beetle larvae extraction by New Caledonian crows (*Corvus moneduloides*)^10^, oyster-cracking by Burmese long-tailed macaques (*Macaca fascicularis aurea*)^7^, and honey-dipping, termite-fishing, and nut-cracking by chimpanzees (*Pan troglodytes spp.*)^8,11,12^. Understanding the extent and causes of individual differences in extractive-foraging tasks is key for identifying variation in factors known to be relevant to evolutionary fitness.

Secondly, there is growing recognition that variation in tool use produces variation in the traces left in the archaeological record, particularly for the signatures produced during percussive behaviours^5^. For example, the number of unintentional flakes produced by wild bearded capuchin monkeys (*Sapajus libidinosus*) whilst cracking nuts with stones will vary according to the tool user, with the skill (or lack thereof) of the individual determining the frequency of mishits and subsequent flakes^13^. Investigating the archaeological signatures of known individuals can inform our understanding of hominin fossil sites and the context behind technological traces^5^. However, there is currently no research on individual-level variation in chimpanzee stone tool use, constituting a significant gap in our understanding of primate technological behaviour and its potential implications for hominin tool use.

Thirdly, identifying systematic variation in technological behaviour is crucial for investigating the factors influencing individual differences in behavioural acquisition via social learning. Indeed, although there are species-typical forms of social learning for particular skills, there is growing evidence that social learning mechanisms vary systematically across individuals^14^. Individual-level variation can develop from ‘individual learning of social learning’, whereby differing experiences with previous social learning opportunities can result in different strategies being employed in the future, or the rate of learning being altered. This phenotypic plasticity can facilitate faster adaptation to environmental changes, potentially having profound effects on evolutionary processes. This is particularly true where model selection and social learning biases guide behavioural acquisition^15–18^. For example, female migrant chimpanzees in the Taï Forest, Côte d’Ivoire, conform to the nut-cracking technique of their new community, even when their previous technique was more efficient (in terms of strikes per nut and foraging speed)^17^. Therefore, assessing variation contributes to our understanding of cultural transmission, and the evolution of complex behavioural traits in primates^19^.

Variation in technological behaviour in nonhuman primates is well established. For example, only male bearded capuchins use sticks as probing tools^20^. In chimpanzees, inter- and intra-population variation in tool use is found across all communities, including differences in technological strategies and efficiency. For example, ant-dipping tool lengths differ between the two neighbouring communities in the Kalinzu Forest, Uganda^21^. Chimpanzees in Loango National Park, Gabon, use different grip types to perforate nests during honey extraction, which is unaffected by the soil hardness^8^. In Gombe National Park, Tanzania, females commence termite-dipping earlier, engage in fishing more frequently, and retrieve more termites per dip than males^22^.

In nut-cracking, females in Taï crack *Coula edulis* and *Panda oleosa* nuts more efficiently than males^23,24^. In contrast, males in Bossou, Guinea, select, use, and transport tools more frequently than females^25^, and spend a significantly greater proportion of their time cracking oil palm (*Elaeis guineensis*) nuts compared to adult females^26^. The number of hits required to successfully open the nutshell was previously found to reach an asymptote in adulthood^27^, and the movement of adult crackers was considered to be stereotyped^28^. However, this finding is constrained to a measure of strikes per nut, which is only one measure of nut-cracking efficiency. Moreover, no research has sought to replicate this finding, and the presence and stability of variation in nut-cracking behaviour has not been investigated using long-term data. The combination of chimpanzees’ long lifespans and the protracted learning periods (3–5 years for nut-cracking^28,29^) required to learn complex tool use necessitates longitudinal studies for assessing developmental trajectories and social learning. Although cross-sectional data can be useful for determining general developmental milestones, they do not allow for tests of individual variation across the lifespan. By obscuring potential variability, this may result in generalisations about population- or species-typical development of a behavioural trait. Tracking the development of a skill across an individual’s life course is key for understanding the ontogenetic processes involved—and potential variation in—fitness-enhancing traits, including extractive technologies.

This study assessed five measures (see Table 1) of post-learning period nut-cracking efficiency for 21 individuals present in a 25-year video archive of wild chimpanzees. Data were gathered from all post-learning period individuals present in the archive for each year they were in the footage (see Figure 2). ‘Post-learning period’ was defined as chimpanzees aged 6 and over, based on the previously established critical learning period lasting until 5-years-old^28^. This research is the first to take such a longitudinal approach to investigate the presence and stability of variation in primate extractive foraging. To our knowledge, this research is also the first to assess the reliability (whether individual differences are captured consistently across measures) and stability (the relationship between measures and whether they hold over time)^1^ of such differences in a wild primate population.

**Figure 2.**
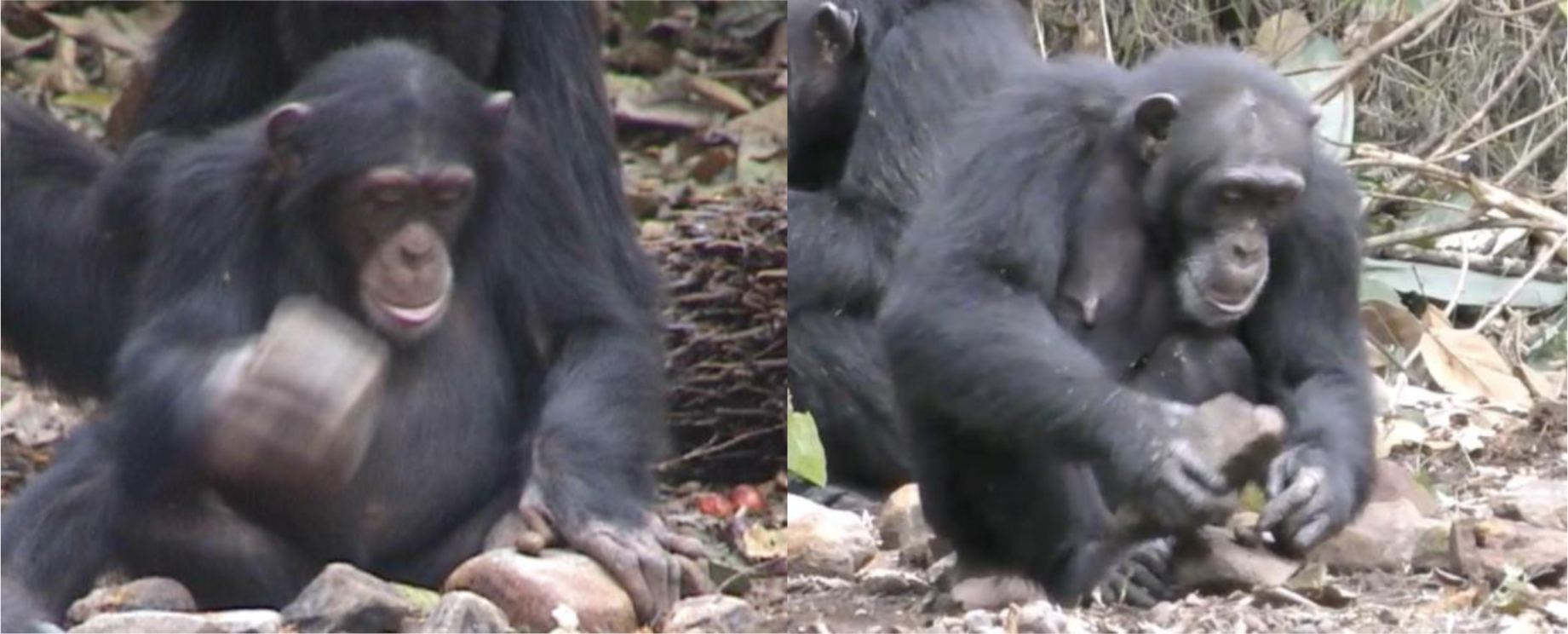
Stills taken from the Bossou video archive. On the left, a six-year-old female chimpanzee, Fanle, cracks an oil palm nut in 2004, the year after finishing her learning period. On the right, the same female chimpanzee cracks an oil palm nut in 2017 at the age of 20. Image credit: Kyoto University.

**Table 1.**
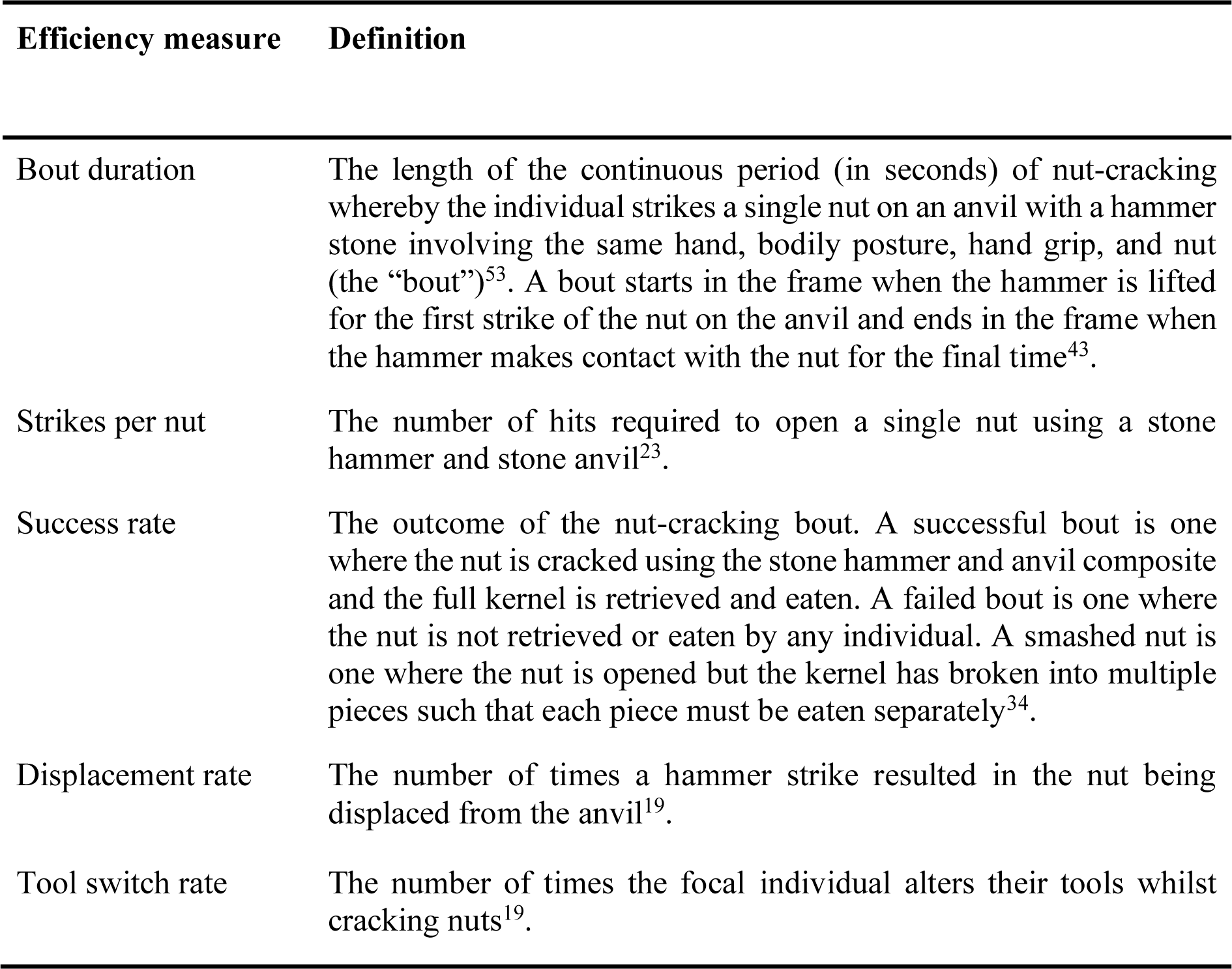
Definitions of efficiency measures.

## Results

### Nut-cracking bouts

A total of 3,882 oil-palm nut-cracking bouts were recorded across 21 chimpanzees (12 female; ages 6–60) over a 25-year period. Of these bouts, 318 (8.17%) ended in failure, 336 (8.66%) ended with a smashed kernel (which results in only partial retrieval of the kernel), and 3,228 (83.17%) ended in the successful retrieval of a whole kernel.

### Individual variation

Analyses (see Tables S2–6) revealed that including random intercepts for individual chimpanzees improved model fit for all measures: *bout duration* (*χ*^2^(1) = 369.49, *p* < 0.0001), *strikes per nut* (*χ*^2^(1) = 475.26, *p* < 0.0001), *success rate* (*χ*^2^(1) = 47.16, *p* < 0.0001), *displacement rate* (*χ*^2^(1) = 87.022, *p* < 0.0001), and *tool switch rate* (*χ*^2^(1) = 4.17, *p* < 0.05). Age had a significant fixed effect on *bout duration* (*t* = 7.724, *p* < 0.001) and *strikes per nut* (*z* = 8.51, *p* < 0.001), *displacement rate* (*z* = -2.40, *p* < 0.05), and *tool switch rate* (*z* = -2.462, *p* < 0.05). There was a significant fixed effect of sex for *tool switch rate* (*z* = -2.502, *p* < 0.05), but not for *bout duration* (*t* = 0.238, *p* > 0.05), *strikes per nut* (*z* = 0.407, *p* > 0.05), *success rate* (*z* = -0.1296, *p* > 0.05), or *displacement rate* (*z* = -1.029, *p* > 0.05).

### Stability of variation

Next, we ranked the random intercepts for each multilevel model; random intercepts represent estimated individual-level effects on the outcome variable in the model. Lower ranks represent greater relative nut cracking efficiency: fewer strikes per nut, shorter bout durations, *etc*. Ranks ranged from 1 to 21 (the total number of individuals in the dataset, see Figure 3). Individual ranks for *bout duration*, *strikes per nut*, *success rate*, and *displacement rate* were moderately-strongly correlated (see Figure S1), meaning that when an individual was ranked highly on one of these measures, they also ranked highly in the other two. However, *tool switch rate* was only weakly–moderately correlated, suggesting it reflects a different underlying construct.

**Figure 3.**
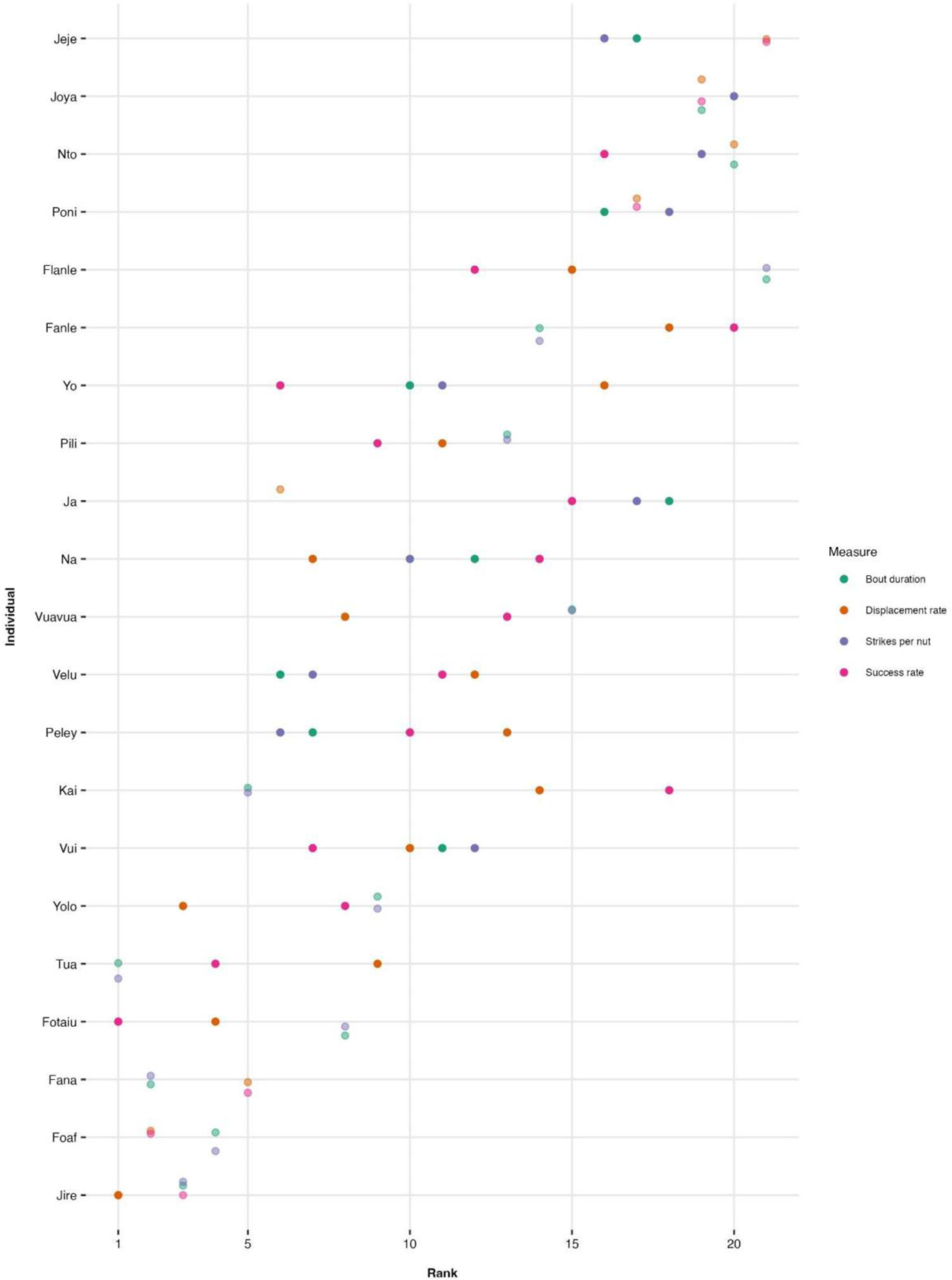
Ranked random intercepts from each multilevel model for the four stable and reliable measures of nut-cracking efficiency.

### Reliability of variation

The consistency of the relative rank of each chimpanzee’s random intercepts obtained in the *bout duration*, *strikes per nut*, *success rate*, and *displacement rate* models indicates reliable individual differences in nut-cracking efficiency. A two-way intra-class correlation (ICC) analysis also indicated good consistency, ICC(C,1) = 0.718, F(20,60) = 11.2, 0.543 < ICC < 0.857, *p* < 0.001. Including *tool switch rate* in the two-way ICC decreased the internal consistency of the efficiency measures (ICC(C,1) = 0.558, F(20,80) = 7.32, 0.366 < ICC < 0.751, *p* < 0.001), further suggesting it reflects a different underlying construct.

### Inter-rater reliability

Two independent, hypothesis-blind coders reviewed 70 hours (8.41%) of footage for inter-coder reliability analyses. Unweighted Cohen’s κ and ICC analyses indicated substantial–excellent agreement and consistency between coders (see Supplementary Information).

## Discussion

This research is the first to establish the presence of long-term stable and reliable systematic individual differences in the technological efficiency of wild chimpanzees. This finding highlights the necessity to move beyond exclusively using averages when investigating daily energy expenditure and activity budgets—factors with large implications for life history and foraging ecology in primates. Variation in time and energy expended for nut-cracking will produce variation in resource allocation to other fitness-enhancing traits. Thus, the extent of variation in nut-cracking efficiency may have large implications for other factors impacting survival and reproduction at the individual-level. In turn, this will inform our understanding of potential individual differences in social learning, which could directly impact the evolutionary fitness of individuals as well as the evolution of cultural traits.

Moreover, this research is the first to demonstrate individual variation in chimpanzee stone tool use. This finding reiterates the need to consider variation in the archaeological signatures left by percussive technological behaviours, with some individuals potentially contributing more to the record than others^5^. Research is now needed to ascertain the sources of this variation. A particular focus on the nut-cracking learning period is key given the known developmental drivers of variation in technological behaviour^22,30,31^.

Each variable measured captured different aspects of ‘efficiency’, with *bout duration*, *strikes per nut*, *success rate*, and *displacement rate* being closely correlated. Bout duration is an indicator of energy intake, as shorter bouts allow more kernels to be retrieved over the course of a feeding session. This may improve an individual’s food security, as they will be at a competitive advantage compared to conspecifics in terms of access to more nuts. Also, longer bouts mean that the individual’s attention is focused on the task for a relatively longer period of time, such that there is reduced time to be allocated to fitness-enhancing behaviours such as defence, socialisation, and grooming. Conversely, the number of strikes required to retrieve the kernel and the success rate of the individual are indicators of energy expenditure relative to energy intake. The number of strikes per nut also dictates how convenient it is for the individual to use stones to open the nuts (rather than, for example, using their teeth or scrounging kernels), with more convenient techniques being favoured due to their increased efficiency^32^. For success rate, future research should investigate why chimpanzees choose to end a nut-cracking bout before the kernel is extracted and assess whether certain individuals stop bouts earlier than others (i.e., do not expend unnecessary energy and are hence more efficient).

Displacement rate measures the ability, or lack thereof, to judge the amount of kinetic energy required to strike a nut. Displacing the nut not only expends more energy than is required, but also elongates the bout duration, particularly if the individual has to travel to retrieve the nut. This also reduces the convenience of using tools. Primate archaeological research should investigate whether individuals with higher displacement rates are also more likely to fracture their stone tools, resulting in potential unintentional flakes^33^.

Our results also suggest that the number of times individuals adjust their tools during a bout reflects a different underlying process. Tool switch rate is a proxy measure for the individual’s ability to select an efficient tool composite and position the stones in such a way to reduce the energetic effort required to extract the kernel. Given the importance of tool properties in determining the efficiency of the behaviour^17,34,35^, learning to select the best tools is also an important aspect of successful foraging though our results suggest that it does not relate to how efficient the individual is at cracking nuts *per se*.

Unlike previous research, we found inconsistent effects of sex on technological efficiency. Sex had no significant effect on bout duration, strikes per nut, success rate, or displacement rate, but there was a significant effect on tool switch rate, with male chimpanzees switching tools less frequently than female chimpanzees. This finding contradicts previous research establishing a female bias in technological behaviour across the genus *Pan*^36^. The lack of sex differences in this research may suggest that the mechanism(s) establishing a female bias in nut-cracking efficiency (measured as the mean number of strikes per nut and nuts per minute) in Taï^24^ are not present in Bossou. However, it may be that the apparent female bias may be an artefact of short-term, cross-sectional analyses, and that the effect does not hold longitudinally. Further long-term research is required to determine whether this finding holds for other populations of nut-cracking chimpanzees.

Age had a significant fixed effect on bout duration, strikes per nut, displacement rate, and tool switch rate. Increasing age corresponds to longer bouts and more strikes per nut, which may reflect greater muscle weakness and the need for more rests in old age. Indeed, there have been many elderly chimpanzees in the Bossou population, with six individuals in the archive being aged 45+. Conversely, increasing age also corresponds to fewer nut displacements and fewer tool switches during bouts. This may reflect older individuals’ greater proficiency at selecting appropriate tools, and better ability at judging the amount of kinetic energy required for each strike. More research is needed to assess the impact of old age on chimpanzee technological capabilities.

The presence of variation in the different measures of nut-cracking efficiency may indicate variation in the underlying cognitive and/or motor capacities. For example, longer bout durations may signify that certain individuals are prone to distraction rather than an individual being inherently less skilled. Therefore, the stable individual variation in nut-cracking efficiency may suggest that there are stable cognitive differences between the individuals, which potentially has implications for performance in other key fitness-enhancing behaviours. Future research should assess whether variation exists in other behaviours in the community to determine the extent of these cognitive differences.

The research is limited by two main points. First, despite the experimental nature of the outdoor laboratory, there are risks of confounding variables. For example, there is no control for what else each individual has consumed in a day or levels of physical activity, both of which could influence motivations for cracking nuts^37^. However, footage selected for analysis was randomly sampled, representing a period of many years, in an attempt to amplify the signal-to-noise ratio.

Second, the outdoor laboratory is a field experimental set up, with nuts and stones being provisioned, which could be argued decreases the validity of the findings. However, the nut-cracking that occurs in the outdoor laboratory is no different from the nut-cracking that the Bossou chimpanzees perform at the natural cracking site of Moblim (07°38’20.7’’ N; 008°30’39.2’’ W)^25^. Moreover, the location of the outdoor laboratory was specifically selected to be on Mont Gban’s summit—the core of the home range for this chimpanzee community— to optimise the frequency of chimpanzees visiting the site^38,39^. This suggests that nut-cracking behaviour remains unchanged regardless of site location, reiterating the ecological validity of the outdoor laboratory.

## Conclusion

This study systematically assessed individual differences in nut-cracking in the Bossou chimpanzees using a long-term video archive of 25 years to longitudinally investigate these differences. Our results establish stable and reliable individual-level differences across four measures of nut-cracking efficiency, shedding light on the underlying cognitive and behavioural diversity in the Bossou chimpanzees. This research contributes to a growing body of evidence finding stable cognitive abilities in great apes, and points to potential variation in the development of this extractive foraging skill. Future research should seek to establish the factors driving this individual variation.

## Methods

### Study site

Bossou is a village in south-eastern Guinea (7°38’71.7’N, 8°29’38.9’W), with a tropical wet seasonal climate and a predominant population of the Manon ethnic group^37,39^. The neighbouring chimpanzee community resides in primary and secondary forest, with a home range of 15 km^2^, although their core area is 7 km^2^. The chimpanzees in Bossou have been studied continuously since 1976, with 21 individuals being present^40^. Since then, the community has been declining in size^41^. The Bossou chimpanzees use a stone hammer-and- anvil composite to extract oil palm nuts^12,33,42^, with individuals requiring complementary coordinated action of both hands to manoeuvre three objects (hammer, anvil, nut) during a nut-cracking bout^43^.

### Study materials

The Bossou chimpanzees have been recorded in every dry season (December–February) since 1988, resulting in a long-term video archive of their behaviour. Researchers observed and videoed the chimpanzees in the ‘outdoor laboratory’^28,38^—a 7 m x 20 m clearing in the core of the community’s home range on Mount Gban (7°38’41.5″ N, 8°29’50.0″ W), which is passed through daily^28,33,38^.

The outdoor laboratory is experimental in nature, with many of the available raw materials (with established weights and dimensions) and nuts being provisioned by the researchers^33^. Observations of behaviour within the outdoor laboratory occur from behind a grass screen along one edge of the clearing. All observation sessions were recorded using at least two standardised camera angles^33^ (wide- and standard-angle lenses), optimising the viewing angles.

### Data collection

Behavioural analysis was conducted using *Behavioural Observation Research Interactive Software* (BORIS, v. 7.11.1)^44^. Of the 1185 videos in the Bossou archive from 1992–2017, 966 videos contained visible nut-cracking bouts by at least one individual, amounting to 832 observation hours. Where possible, the videos from the standard-angle lens data were used for analysis. If behaviour was obscured in this footage, the equivalent video with the wide-angle lens was uploaded to attempt to observe the behaviour. If the behaviour remained unobservable, the bout(s) were excluded. No footage was collected in 2001 or 2011, and so these are absent from the data set.

Data from each year each focal individual was present in the archive was collected. Multiple bouts (up to 20) per individual per year were recorded to establish the degree of within-individual variation in efficiency, while also producing more independent data points, allowing between-individual variation to be assessed. The full data collection protocol can be found in the Supplementary Information.

Ethical approval and permissions to conduct scientific research in the Bossou community were obtained by each contributor to the video archive from the Direction Générale de la Recherche Scientifique et de l’Innovation Technologique (DGERSIT) and the Institut de Recherche Environnementale de Bossou (IREB) in Guinea. The video footage was used with the permission of Tetsuro Matsuzawa and Kyoto University, Japan. No further ethical approval was required by the School of Anthropology and Museum Ethnography, University of Oxford, or the Social Science Division, University of Oxford.

### Statistical analysis

All bouts where an infant was clinging to the focal subject were removed from analyses (*n* = 210), as this was exclusive to certain female chimpanzees (Fana, Fanle, Fotaiu, Jire, Pili, Velu, Vuavua, and Yo) and so could reduce the internal validity of the findings by altering the efficiency of these individuals. Removing these bouts ensured that all subjects were compared under equal circumstances. This left 3,672 bouts for analysis.

Only data for bouts in which a kernel was retrieved were included in the analyses for *bout duration* and *strikes per nut* (*n* = 3,367). Excluding the ‘Failed’ bouts here ensured that the amount of time it took to access the energetic reward of the enclosed kernel was analysed. The full data set (*n* = 3,672) was analysed for *success rate*, *displacement rate*, and *tool switch rate*.

All analyses were performed using R (v. 4.3.2)^45^ and RStudio (v. 2023.09.1+494) for MacOS. The significance level (*α*) was 0.05 for all analyses. Multilevel models were constructed to test for individual differences in the five measures of efficiency. Individual chimpanzees comprised the level-two random factor, while age and sex were included as fixed effects. Simple models (without random effects) were constructed and compared against the multilevel models (see Tables S2–S6); ANOVAs were used to determine the model with the best fit. The ANOVAs (two-tailed) assessed whether including a random intercept for individual chimpanzees significantly decreased the model prediction error. The model with the smallest prediction error (AIC and -2-log-likelihood values) was selected for each of the five components of efficiency. The linear mixed-effects model for *bout duration* was fitted using the *lme4*^46^ and *lmerTest*^47^ packages. Initially, a simple linear multilevel model (subject as a random intercept) was constructed, with age and sex as fixed effects. However, this model fit the data poorly, and so the normality of the outcome variable *bout duration* was assessed using a histogram. The data were found to be strongly right skewed, and so they were log transformed to normalise the distribution and correct for heteroskedasticity^48^.

Given that the count data for *strikes per nut* did not contain zeros, zero-truncated poisson generalised linear (mixed) models (GL(M)Ms) were fitted using the truncated_poisson family in the *glmmTMB* package^49^, with age and sex as fixed effects. However, the check_dispersion function from the *performance* package^50^ detected overdispersion in the multilevel model (subject as a random intercept). As such, zero-truncated negative binomial GL(M)Ms were fitted with quadratic parameterisation.

As the data for *success rate* were ordinal, cumulative link models (CLMs) and cumulative link mixed models (CLMMs) were fitted using the *ordinal* package^51^. The ‘outcome’ factor was ordered to Failed, Smash, Successful, to account for the increasing degrees of efficiency with each increasing level of outcome^34^. The Hessian matrices were calculated for all models to allow the model summaries to be called. The number of quadrature points used in the adaptive Gauss-Hermite quadrature approximation was set to 7. Age was originally included as a fixed effect but was removed as the model did not converge. Sex remained as a fixed effect.

Negative binomial GL(M)Ms were fitted for *displacement rate* and *tool switch rate* using the *glmmTMB* package. All models specified zero-inflation as being equal for all observations, given the large number of zeros in the data where an individual did not displace the nut (*n* = 2909) or switch their tools (*n* = 3308) in a bout.

The individuals’ random intercepts were ranked for the *bout duration*, *strikes per nut*, *success rate*, *displacement rate*, and *tool switch rate* models, and a correlation matrix was constructed to determine whether they represent the same underlying construct. The two-way ICC used to assess the stability of individual variation was selected according to recommended guidelines^52^.

## Data availability

A minimum dataset can be found in the following public repository: https://github.com/arranjdavis/chimpanzee_nut_cracking_efficiency

## Code availability

All analysis code needed to reproduce the results and figures reported in the paper and the Supplementary Information can be found in the following public repository: https://github.com/arranjdavis/chimpanzee_nut_cracking_efficiency

## Acknowledgements

We thank the Direction Générale de la Recherche Scientifique et de l’Innovation Technologique (DGERSIT) and the Institut de Recherche Environnementale de Bossou (IREB) in Guinea for research authorisation. The video data for this research was provided by Tetsuro Matsuzawa and Kyoto University, Japan. The original video archive was digitised, organised, and systematised by Daniel Schofield, Department of Engineering Science, University of Oxford. Special thanks to all the field researchers and assistants at Bossou, specifically Henry Didier Camara, Cé Vincent Mami, Gouanou Zogbila, Boniface Zogbila, Jules Doré, Guana Goumy, Tino Camara, Pascal Goumy, Paquílé Cherif, Jiles Gondo Doré, and Marcel Doré. Thank you to all the researchers who contributed to the Bossou archive since its inception. Thank you to Alexander Mielke for help cataloguing the Bossou video archive. Thank you to Dora Biro and Katarina Almeida Warren for feedback. Thank you to Zhangzhuoran Dai and Victoire Martignac for being independent, hypothesis-blind coders for the inter-rater reliability analyses.

## Author contributions

**S.B.** conceived of the study, designed, and coordinated the study, collected data, participated in the analysis and visualisation of the data, and wrote the manuscript. **E.C.** provided supervision, participated in the design of the study, and commented on the manuscript. **A.J.D.** analysed and visualised the data and commented on the manuscript. **T.M.** collected the original data set and commented on the manuscript. **S.C.** collected the original data set, provided supervision, participated in the design of the study, and commented on the manuscript.

## Competing interests

The authors declare no competing interests.

**Supplementary information** is available for this paper.

**Correspondence and requests for materials** should be addressed to Sophie Berdugo.

