## Supplementary Information for "Stable long-term individual variation in chimpanzee technological efficiency"

### Data collection protocol

First, all available footage was systematically reviewed to ascertain the party composition in each video, and verify which chimpanzees were visible cracking nuts in the footage. Following this, the videos' unique identifiers (UIDs) for each year were placed into ascending order and allocated an integer code starting from one, incrementally increasing by one. Vectors for each focal individual in each year were constructed in RMarkdown[1], each comprising the code for each video the individual was nut-cracking. Each vector was sampled without replacement to create a random order of video codes for all individuals. A seed was set to make the sequence of random codes replicable.

From this process, it became apparent that some individuals in the community (for example, Velu and Fana) were present and nut-cracking in the footage considerably less frequently than the other community members. To reduce potential bias introduced from the varying sample sizes for each subject, data from all nut-cracking bouts for the rare individuals (defined as being present and having observable nut-cracking bouts in ≤ 25% of videos for a given year) were collected. Where other chimpanzees had observable nut-cracking bouts in this footage, data from their bouts were also collected. This allowed for the effects of seasonality on nut hardness to be partially controlled. Thereafter, the videos were selected from the randomly ordered vectors (present and nut-cracking in > 25% of videos for a given year), starting from the least common of the remaining chimpanzees. This process continued until at least 20 nut-cracking bouts had been recorded for each individual.

Data from each year each individual was present in the archive was collected. Multiple bouts per individual per year were recorded to establish the degree of within-individual variation in efficiency, while also producing more independent data points, allowing between-individual variation to be assessed. This reduced the sampling error and random variation found between years, and hence amplified the signal-to-noise ratio. This was to ensure that the data collected were reliable, and representative of the true behaviour of the group.

Lastly, to ensure the measures of efficiency were recorded accurately, only bouts which were clearly visible (i.e., observable) were coded. Visible bouts were those where the focal individual was facing the camera and the nut, hammer, and anvil could be seen, and those where the individual was not directly facing the camera, but the nut, hammer, and anvil could be seen. At the end of each bout, whether or not the complete bout was observed was recorded. Incomplete bouts were removed prior to analysis. This reduced the risk of systematic bias being introduced into the sampling procedure by the recording period ending prior to the termination of the behaviour, or because the focal subject became occluded[2].

**Subject information**

***Table S1.*** Focal subject information, with the years that they were observably cracking nuts in the Bossou archive during their post-learning period.

| Subject | Sex | Nut-cracking hand | Observation years | Age (years) | Bouts observed |
|---|---|---|---|---|---|
| Fana | F | Right* | 1992–2017 | 36–60 | 386 |
| Jire | F | Left | 1992–2017 | 34–59 | 346 |
| Yo | F | Left | 1992–2016 | 32–56 | 347 |
| Tua | M | Left | 1992–2012 | 35–55 | 338 |
| Velu | F | Right | 1992–2015 | 33–55 | 248 |
| Kai | F | Right | 1992–2002 | 42–52 | 164 |
| Foaf | M | Right | 1992–2017 | 12–37 | 416 |
| Fanle | F | Right | 2004–2017 | 6–20 | 234 |
| Jeje | M | Left | 2004–2017 | 7–20 | 236 |
| Yolo | M | Left | 1998–2009 | 6–17 | 208 |
| Peley | M | Left | 2005–2012 | 6–14 | 133 |
| Pili | F | Right | 1993–2000 | 6–13 | 102 |
| Vui | M | Left | 1992–1999 | 6–13 | 123 |
| Vuavua | F | Left | 1998–2004 | 6–12 | 106 |
| Fotaiu | F | Right | 1998–2003 | 6–11 | 90 |
| Na | M | Right | 1992–1996 | 7–11 | 82 |
| Ja | F | Right | 1992–1993 | 9–10 | 22 |
| Poni | M | Right | 2000–2002 | 7–9 | 41 |
| Joya | F | Left | 2010–2012 | 6–8 | 25 |
| Flanle | M | Left | 2014 | 6 | 14 |
| Nto | F | Right | 2000 | 6 | 11 |

*Note*: * = switched to her right hand after her left arm became paralysed in 1996.

### Model outputs

***Table S2.*** Simple and multilevel model outputs for log bout duration efficiency measure.

| | **Log bout duration** | | | **Log bout duration** | | |
|---|---|---|---|---|---|---|
| *Predictors* | *Estimates* | *CI* | *p* | *Estimates* | *CI* | *p* |
| (Intercept) | 1.21 | 1.11–1.30 | **<0.001** | 0.59 | 0.18–1.01 | **0.005** |
| Age | -0.01 | -0.01–-0.00 | **<0.001** | 0.02 | 0.02–0.03 | **<0.001** |
| Sex [Male] | -0.35 | -0.42–-0.27 | **<0.001** | 0.07 | -0.53–0.67 | 0.812 |
| **Random Effects** | | | | | | |
| $\sigma^2$ | | | | 0.94 | | |
| $\tau_{00}$ | | | | 0.46 Subject | | |
| ICC | | | | 0.33 | | |
| N | | | | 21 Subject | | |
| Observations | 3367 | | | 3367 | | |
| $R^2$ / $R^2$ adjusted | 0.023 / 0.022 | | | 0.086 / 0.387 | | |

*Note*: The confidence intervals are calculated using the standard error for the fixed effects. The random effects residual variance ($\sigma^2$) and intercept variance ($\tau_{00}$) are presented.

***Table S3.*** Zero-truncated negative binomial simple and multilevel model outputs for strikes per nut efficiency measure.

| | **Strikes per nut** | | | **Strikes per nut** | | |
|---|---|---|---|---|---|---|
| *Predictors* | *Incidence Rate Ratios* | *CI* | *p* | *Incidence Rate Ratios* | *CI* | *p* |
| (Intercept) | 3.04 | 2.76–3.34 | **<0.001** | 1.68 | 1.14–2.47 | **0.008** |
| age | 1.00 | 1.00–1.00 | 0.268 | 1.02 | 1.02–1.03 | **<0.001** |
| sex [Male] | 0.71 | 0.66–0.77 | **<0.001** | 1.12 | 0.65–1.94 | 0.684 |
| **Random Effects** | | | | | | |
| $\sigma^2$ | | | | 0.49 | | |
| $\tau_{00}$ | | | | $0.39_{\text{Subject}}$ | | |
| ICC | | | | 0.44 | | |
| N | | | | $21_{\text{Subject}}$ | | |
| Observations | 3367 | | | 3367 | | |
| $R^2$ conditional / $R^2$ marginal | NA / 0.028 | | | 0.150 / 0.525 | | |

*Note*: The confidence intervals are calculated using the standard error for the fixed effects. The random effects residual variance ($\sigma^2$) and intercept variance ($\tau_{00}$) are presented.

***Table S4.*** Cumulative link simple and multilevel model outputs for success rate efficiency measure.

| | **Success rate** | | | **Success rate** | | |
|---|---|---|---|---|---|---|
| *Predictors* | *Odds Ratios* | *CI* | *p* | *Odds Ratios* | *CI* | *p* |
| Failed\|Smash | 0.09 | 0.08–0.10 | **<0.001** | 0.09 | 0.07–0.12 | **<0.001** |
| Smash\|Successful | 0.20 | 0.18–0.23 | **<0.001** | 0.21 | 0.17–0.27 | **<0.001** |
| Sex [Male] | 0.96 | 0.81–1.14 | 0.641 | 0.98 | 0.67–1.43 | 0.897 |
| N | | | | 21 $_{\text{Subject}}$ | | |
| Observations | 3672 | | | 3672 | | |
| $R^2$ Nagelkerke | 0.000 | | | NA | | |

***Table S5.*** Simple and multilevel zero-inflated negative binomial model outputs for displacement rate efficiency measure.

| | **Displacement rate** | | | **Displacement rate** | | |
|---|---|---|---|---|---|---|
| *Predictors* | *Incidence Rate Ratios* | *CI* | *p* | *Incidence Rate Ratios* | *CI* | *p* |
| (Intercept) | 0.56 | 0.47–0.65 | **<0.001** | 0.43 | 0.30–0.62 | **<0.001** |
| (Intercept) | 0.56 | 0.47–0.65 | **<0.001** | 1.51 | 1.38–1.69 | |
| age | 0.98 | 0.98–0.98 | **<0.001** | 0.99 | 0.98–1.00 | **0.016** |
| sex [Male] | 0.79 | 0.68–0.92 | **0.002** | 0.81 | 0.54–1.21 | 0.303 |
| (Intercept) | 1.60 | 1.45–1.80 | | 0.43 | 0.30–0.62 | **<0.001** |
| (Intercept) | 1.60 | 1.45–1.80 | | 1.51 | 1.38–1.69 | |
| **Zero-Inflated Model** | | | | | | |
| (Intercept) | 0.00 | 0.00–Inf | 0.994 | 0.00 | 0.00–Inf | 0.993 |
| **Random Effects** | | | | | | |
| $\sigma^2$ | | | | 1.68 | | |
| $\tau_{00}$ | | | | 0.16 Subject | | |
| ICC | | | | 0.09 | | |
| N | | | | 21 Subject | | |
| Observations | 3672 | | | 3672 | | |
| $R^2$ conditional / $R^2$ marginal | NA / 0.056 | | | 0.020 / 0.104 | | |

*Note*: The confidence intervals are calculated using the standard error for the fixed effects. The random effects residual variance ($\sigma^2$) and intercept variance ($\tau_{00}$) are presented.

***Table S6.*** Simple and multilevel zero-inflated negative binomial model outputs for tool switch rate efficiency measure.

| | **Tool switch rate** | | | **Tool switch rate** | | |
|---|---|---|---|---|---|---|
| *Predictors* | *Incidence Rate Ratios* | *CI* | *p* | *Incidence Rate Ratios* | *CI* | *p* |
| (Intercept) | 0.19 | 0.15–0.24 | **<0.001** | 0.18 | 0.14–0.25 | **<0.001** |
| (Intercept) | 0.19 | 0.15–0.24 | **<0.001** | 1.32 | 1.21–1.50 | |
| age | 0.99 | 0.98–1.00 | **0.002** | 0.99 | 0.98–1.00 | **0.014** |
| sex [Male] | 0.66 | 0.53–0.83 | **<0.001** | 0.68 | 0.50–0.92 | **0.012** |
| (Intercept) | 1.33 | 1.22–1.51 | | 0.18 | 0.14–0.25 | **<0.001** |
| (Intercept) | 1.33 | 1.22–1.51 | | 1.32 | 1.21–1.50 | |
| **Zero-Inflated Model** | | | | | | |
| (Intercept) | 0.00 | 0.00–Inf | 0.995 | 0.00 | 0.00–Inf | 0.995 |
| **Random Effects** | | | | | | |
| $\sigma^2$ | | | | 2.43 | | |
| $\tau_{00}$ | | | | 0.04 Subject | | |
| ICC | | | | 0.02 | | |
| N | | | | 21 Subject | | |
| Observations | 3672 | | | 3672 | | |
| $R^2$ conditional / $R^2$ marginal | NA / 0.017 | | | 0.016 / 0.031 | | |

*Note*: The confidence intervals are calculated using the standard error for the fixed effects. The random effects residual variance ($\sigma^2$) and intercept variance ($\tau_{00}$) are presented.

### Inter-rater reliability

Two independent, hypothesis-blind coders were recruited to test the between-observer reliability of the five efficiency components. This took place following the pilot research to ensure that 1) the coding scheme was finalised prior to the main data collection, and 2) that the coding scheme was consistent throughout the investigation, as any ambiguities in the behavioural category definitions were clarified *a priori*. This was important for reducing potential disagreement, and hence increasing reliability, between coders.

Both independent coders received thorough training for using the coding scheme and the BORIS software. Thereafter there was no consultation between coders, although the identity of the individuals in the videos were provided to assist with the accuracy of the behavioural coding.

The videos for reliability analysis were randomly selected to reduce the risk of bias. A combined total of 70 hours of observation was completed by the independent coders.

Cohen's κ was calculated to determine the extent of agreement between coders for *success rate* as the measure was categorical. All statements of the strength of the agreement between the coders are in accordance with standardised benchmarks[3].

Numerical data were compared using intraclass correlations (ICCs). Here, two-way random-effects models were used. The type was selected to be ‘single rater’ since measurements were not averaged across the *k* number of raters. Finally, ‘definition’ varies depending on the variable. *Strikes per nut*, *displacement rate*, and *tool switch rate* were selected as ‘absolute agreement’ to check if scores matched exactly across coders. *Bout duration* was selected as ‘consistency’ to determine the extent to allow for systematic error[4]. All statements of the strength of the agreement between the coders are in accordance with standardised guidelines[4]. Analyses were performed using the *irr* package[5]. ICC scores to assess the absolute agreement between the three raters can be found in Table S7. For *bout outcome*, the agreement between the three coders was substantial, κ = 0.771, and greater than what would be expected by chance, $Z$ = 19.5, $p$ <0.0001. For *bout duration*, a single-rater, consistency, two-way model ICC analysis found ICC(C,1) = 0.991, F(424,424) = 214, 0.989 < ICC < 0.992, p < 0.000, indicating excellent consistency.

***Table S7.*** ICC calculations using single-rating, absolute agreement, two-way random-effects models.

| | | 95% Confidence Interval | | F Test with True Value 0 | | | |
|---|---|---|---|---|---|---|---|
| | ICC | Lower bound | Upper bound | Value | *df1* | *df*2 | Sig. |
| Strikes per nut | 0.986 | 0.984 | 0.989 | 147 | 424 | 424 | **<0.001** |
| Displacement rate | 0.893 | 0.872 | 0.91 | 17.6 | 424 | 424 | **<0.001** |
| Tool switch rate | 0.708 | 0.652 | 0.755 | 6.03 | 424 | 305 | **<0.001** |

### Stability of efficiency measures

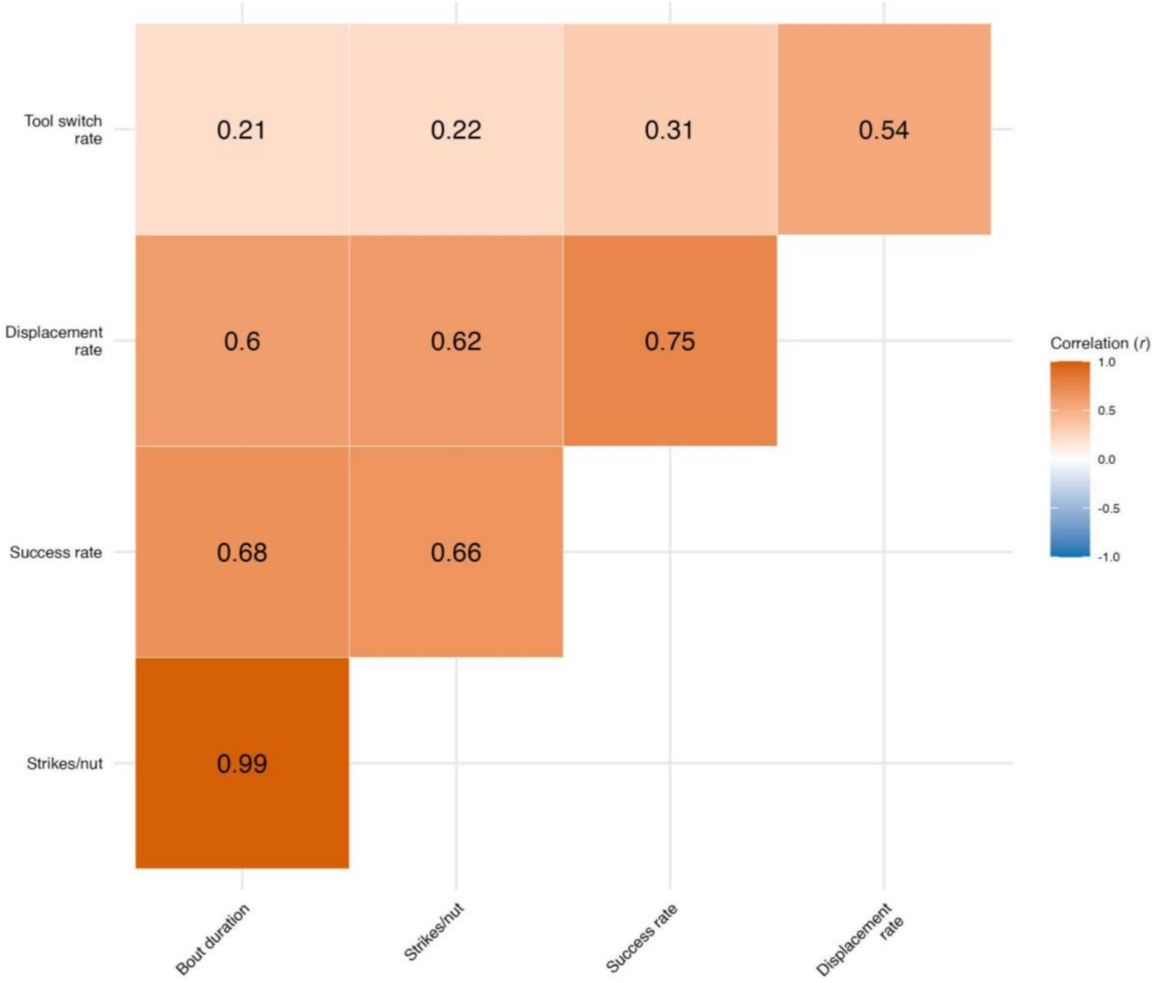


***Figure S1.*** Correlation matrix for the Pearson’s *r* correlation coefficient between the rankings for all pairs of nut-cracking efficiency measures.

### Assumption checks

After fitting the models, the assumptions were checked to ensure inferences could be drawn from the findings. Multicollinearity between predictor variables for the *bout duration* model was checked using the *vif* function (variance inflation factor; VIF) in the *car* package[6]. The VIF was 1.13, indicating no issues of multicollinearity.

For the linear mixed-effects model (*bout duration*), the normality of the residuals was assessed visually using QQ-plots and homoscedasticity was checked by plotting the fitted values against the squared residuals[7]. For the cumulative link model (*success rate*), surrogate residuals[8] were obtained using the *sure* package[9]. We performed assumption checks on the single-level model (CLM) as the package does not currently support multilevel models (CLMM). We assumed this would be sufficient as only the intercepts were allowed to vary in the CLMM. The normality of the surrogate residuals was assessed visually using a QQ-plot and homoscedasticity was checked by plotting the fitted values against the surrogate residuals[10]. For all models, the normality of the random intercepts were assessed using QQ-plots and Shapiro-Wilk tests. Results indicated no significant deviations from normality.

We evaluated the multilevel models using influence diagnostics from the *influence.ME* package[11]. DFBETA values were calculated for each model to assess whether any individuals (i.e., the level two parameter) had an outsized influence on the results of the models. For *bout duration,* two individuals (Fana and Yo) had DFBETA values above the $2/\sqrt{n}$ cut-off[12], indicating that their data were influential. We re-ran the multilevel model (with individual as a random intercept, and age and sex as fixed effects) excluding their data and compared it to a simple linear model (without random effects). The random intercept model fit the data significantly better than the simple model ($\chi^2(1) = 203.75$, $p < 0.0001$), and as such we kept their data included in the model. We found no influential individuals in the *strikes per nut, displacement rate,* and *tool switch rate* models.
